# variant2literature: full text literature search for genetic variants

**DOI:** 10.1101/583450

**Authors:** Yin-Hung Lin, Yu-Chen Lu, Ting-Fu Chen, Jacob Shujui Hsu, Ko-Han Lee, Yi-Wei Cheng, Yi-Chieh Chen, Jhih-Sheng Fan, Chien-Ta Tu, Chen-Ming Hsu, Chih-Chen Chou, Pei-Lung Chen, Yi-Chin Ethan Tu, Chien-Yu Chen

## Abstract

**Motivation:** Whole genome sequencing (WGS) by next-generation sequencing produces millions of variants for an individual. The retrieval of biomedical literature for such a large number of genetic variants remains challenging, because in many cases the variants are only present in tables as images, or in the supplementary documents of which the file formats are diverse.

**Results:** The proposed tool named variant2literature from the TaiGenomics (Toolkits for AI genomics) resolves the problem by incorporating text recognition with image processing. In addition to the adoption of advanced image-based text retrieval, the recall rate of finding the literature containing the variants of interest is further improved by employing the skill of variant normalization. Different variant presentations are transformed into chromosome coordinates (standard VCF format) such that false negatives can be largely avoided. variant2literature is available in two ways. First, a web-based interface is provided to search all the literature in PMC Open Access Subset. Second, the command-line executable can be downloaded such that the users are free to search all the files in a specified directory locally.

**Availability:** http://variant2literature.taigenomics.com/

## 1 Introduction

Whole-exome sequencing (WES) and whole-genome sequencing (WGS) become more and more popular in diagnosis of diseases and discovery of phenotype-related variants [1–3]. The annotation of variants involves a critical step that finds literature to explain the potential influence of a discovered variant on protein functions, cell behaviors, organ normality, etc. In this regard, many efforts have been made to improve the search performance [4, 5]. Even though, we observed that the sensitivity of variant queries still has space to increase. For example, when a researcher sends a query in the literature database, it is most likely that the related papers are missing in the result page owing to the fact that the search is not comprehensively performed on the full text of the papers as a well as all the supplementary files. The challenge of this task is that most accessible literature are released in PDF format. In this study, we used PubMed Central (PMC) Open Access Subset (PMC OA) to demonstrate that the adoption of image processing followed by named-entity recognition (NER) and variant normalization can largely increase the recall rate of literature search.

In addition to tackle the barrier due to diverse file formats, variant2literature also resolves the format issue of variant queries by translating all types of variant presentations into chromosome-based coordinates. The literature files were preprocessed to record all the variants using chromosome-based coordinates in a local database. Moreover, if a gene has multiple isoforms, a known variant of one particular isoform will be also annotated on the other isoforms with corrected coordinates. Meanwhile, RSID (Reference SNP cluster ID) is also acceptable. All of these designs were shown to improve the sensitivity of variant query on literature search. Currently, variant2literature accepts documents files in XML, PDF, DOCX, DOC, XLSX, and CSV formats. Both web interface and command-line executable are available now.

## 2 Materials and methods

When handling PDF files, fasterRCNN [6] was used to detect table regions. Recognition of table structures was performed by in-house rule-based methods. Figure and table captions in PDF were retrieved by PDFFigures 2.0 [7]. variant2lieterature used tmVar [8] to extract variants from paragraphs and tables, and used GNormplus [9] to extract genes from paragraphs and tables. It is important that variant representations were made consistent, in-cluding the variants in the literature and the user queries. For example, “GJB2, c.109G>A”, “GJB2, p.Val37Ile”, “GJB2, V37I”, and “rs72474224” all represent the same variant. Moreover, “TNNT2, R141W”, “TNNT2, R134W”, and “TNNT2, R143W” might present the same variant owing to position shifts on different isoforms of TNNT2. After normalization, the variants and the associated genes were recorded in VCF format [10]. Accordingly, the literature files were indexed by the normalized variants.

variant2literature is available in two ways: web interface and command-line executable. In the web server, query can be made in different ways. Some examples can be found on the web site (http://variant2literature.taigenomics.com/). Meanwhile, uploading a VCF file is acceptable in the web interface.

## 3 Results and discussions

The disease ‘deafness’ is used as an example to demonstrate the power of variant2literature. The Deafness Variation Database (DVD) [11] contains 876,135 known variants (v8.1.2017-11-08). There are 10,036 variants being linked to 3,095 papers, constituting 18,410 variant-paper pairs. Among the 3,095 papers, 206 papers can be found in the PMC OA database, containing 962 variant-paper pairs, involving 874 variants. It is noticed that not all the variant-paper relationships for these 874 variants were curated in the DVD database. In this regard, three domain experts were recruited in this study to identify the missing variant-paper pairs present in the 206 papers for these 874 variants. Finally, 996 pairs were recorded as true relationships.

After invoking variant2literature on 874 variants against the 206 pa-pers, 1,016 pairs were reported as positives, resulting a recall rate of 98.39% and a precision rate of 96.46%. For comparison, we performed the same query of variant set on LitVar [4]. By uploading the 874 variants to LitVar, 672 pairs were reported as positives, resulting a recall rate of 40.15% and a precision rate of 96.43%.

**Table 1.** Performance of variant2literature with different settings.

| Methods | Variant-paper pairs predicted | Recall | Precision | F1 score |
| --- | --- | --- | --- | --- |
| variant2literature (without suppl.) | 444 | 43.47% | 97.52% | 0.60 |
| variant2literature (with suppl.) | 831 | 80.42% | 96.39% | 0.88 |
| variant2literature (with suppl. + table text retrieval) | 1,016 | 98.39% | 96.46% | 0.97 |
| LitVar | 672 | 40.15% | 96.43% | 0.57 |

Abbreviation: suppl. stands for supplementary files.

### 3.1 Limitations

The following limitations of variant2literature should be noticed. First, when a user specifies a query like this: ‘Gene_name, A123C’, variant2literature cannot tell whether the position 123 is the coordinate of bases on cDNA or residues on proteins. Currently, variant2literautre assumes the number is regarding residue coordinates. Second, variant2literature considers all potential isoform coordinates of a gene when searching variants in literature, this might result in some false positives. Such limitations will be gradually resolved according to user feedback in the future.

## Acknowledgements

The authors would like to thank Ministry of Science and Technology (MOST), R. O. C., for the financial support under the contract: MOST 105-2221-E-002-129-MY3. The funder had no role in the design, collection, analysis, or interpretation of the data; writing the manuscript; or the decision to submit the manuscript for publication.

